# Computational modeling of C-terminal tails to predict the calcium-dependent secretion of ER resident proteins

**DOI:** 10.1101/2021.03.21.435734

**Authors:** Kathleen A. Trychta, Bing Xie, Ravi Kumar Verma, Min Xu, Lei Shi, Brandon K. Harvey

## Abstract

The lumen of the endoplasmic reticulum (ER) has resident proteins that are critical to perform the various tasks of the ER such as protein maturation and lipid metabolism. These ER resident proteins typically have a carboxy-terminal ER retention sequence (ERS). The canonical ERS is Lys-Asp-Glu-Leu (KDEL) and when an ER resident protein moves from the ER to the Golgi, KDEL receptors (KDELRs) in the Golgi recognize the ERS and return the protein to the ER lumen. Depletion of ER calcium leads to the mass departure of ER resident proteins in a process termed exodosis, which is also regulated by KDELRs. Here, by combining computational prediction with machine learning-based models and experimental validation, we identify carboxy tail sequences of ER resident proteins divergent from the canonical “KDEL” ERS. Using molecular modeling and simulations, we demonstrated that two representative non-canonical ERS can stably bind to the KDELR. Collectively, we developed a method to predict whether a carboxy-terminal sequence acts as a putative ERS that would undergo secretion in response to ER calcium depletion and interact with the KDELRs. Identification of proteins that undergo exodosis will further our understanding of changes in ER proteostasis under physiological and pathological conditions where ER calcium is depleted.

## Introduction

The classical secretory pathway involves the movement of proteins from the endoplasmic reticulum (ER) to the Golgi and then to the cell surface. Resident proteins of the ER counteract this flow by using short peptide sequences. For soluble proteins, a carboxy-terminal (C-terminal) ER retention sequence (ERS) interacts with KDEL receptors in the Golgi membrane to return proteins back to the ER lumen in a COPI-mediated manner (Munro and Pelham, 1987, Lewis and Pelham, 1992a, Orci et al., 1997). Mammals express three KDEL receptor isoforms, KDELR1, KDELR2, and KDELR3, all of which function as part of a Golgi-to-ER retrieval pathway to maintain the ER proteome (Lewis and Pelham, 1990, Lewis and Pelham, 1992b, Hsu et al., 1992, Collins et al., 2004, Raykhel et al., 2007). Maintaining ERS-containing proteins within the ER lumen is necessary for these proteins to carry out the diverse functions of the ER including protein trafficking and modification and lipid and carbohydrate metabolism. Loss of these proteins from the ER lumen may compromise ER function and lead to gain of function elsewhere inside or outside of the cell.

In mammals, the KDEL (lysine-aspartate-glutamate-leucine) motif is considered the canonical ERS. The first observations of a luminal ER retention sequence noted that three ER resident proteins (BiP/Grp78, Grp94, protein disulfide isomerase) shared a C-terminal KDEL sequence and that deletion of this sequence resulted in secretion of the protein (Munro and Pelham, 1987). Further, appending this KDEL sequence to a secretory protein resulted in a failure of secretion and protein accumulation within the ER (Munro and Pelham, 1987). Subsequent studies identified KEEL, RDEL, and QEDL as putative ERS (Mazzarella et al., 1990, Mazzarella et al., 1994, Fliegel et al., 1990). A bioinformatic approach intended to identify ER localization motifs found 46 putative ERS even while using a data set restricted to human proteins with the terminal four amino acids *XX*[DE][FLM] (Raykhel et al., 2007). As the name implies, the current understanding of ER retention sequences focuses on their ability to retain proteins in the ER. However, recent findings suggest that an important functional significance of ERS lies in their ability to regulate protein secretion. While Raykhel et al. has already provided an analysis of C-terminal motifs that localize proteins to the ER, no analysis has yet been reported that systematically investigates the connection between C-terminal motifs and secretion. This definition of what constitutes an ERS was further expanded upon by Alanen et al. showing that the -5 and -6 amino acids also play a role in determining the ER localization of a protein (Alanen et al., 2011). In fact, expanding to five or six residues for some protein tails caused ER localization whereas the four amino acids did not (Alanen et al., 2011). Thus, the amino acids of the carboxy terminus that determine ER retention can extend beyond the final four C-terminal residues of the protein.

ERS play a role in regulating the secretion of proteins as well. A study of four ER luminal proteins with a C-terminal KDEL (BiP/Grp78, Grp94, protein disulfide isomerase) showed that these proteins were secreted from the cell following treatment with calcium ionophores suggesting that ERS-containing proteins are sensitive to changes in intracellular calcium (Booth and Koch, 1989). More recent studies on ER resident esterases and the ER resident protein mesencephalic astrocyte-derived neurotrophic factor (MANF) demonstrated that the secretion of these proteins is increased specifically following ER calcium depletion (Henderson et al., 2013, Glembotski et al., 2012, Trychta et al., 2018b). Furthermore, in several diseases ERS-containing proteins have been found in extracellular fluid (e.g. serum, plasma, saliva, cerebrospinal fluid) and their presence may be a symptom of disease (Supplementary Table S1). In fact, ER calcium depletion triggered a mass redistribution of ERS proteins from the ER lumen to the extracellular space, a process termed ER exodosis, which is dependent on the level of KDEL receptors (Trychta et al., 2018a). Other triggers of ER exodosis, such as ischemia, have also been identified (Trychta et al., 2018a). A comparison of >100 putative ERS appended to the C-terminus of *Gaussia* luciferase (GLuc) protein revealed varied responses to ER calcium depletion with some ERS being more readily secreted in response to ER calcium depletion than others (Trychta et al., 2018a). This raises the possibility that ERS-containing proteins could act as biomarkers of exodosis in a variety of disease states. The ability to predict whether an ERS confers secretion following ER calcium depletion would be useful for understanding how pathological states affect cellular proteostasis and how mutations may affect trafficking of ERS-containing proteins.

## Methods

### Plasmid construction

GLuc-ERS proteins were created as previously described resulting in pLenti6.3-MANF sigpep-GLuc-ERS constructs (Trychta et al., 2018a). Briefly, custom forward and reverse oligonucleotides coding for each specific seven amino C-terminal tails were synthesized (Integrated DNA Technologies), formed into an oligonucleotide duplex, then inserted into a pLenti6.3-MANF sigpep-GLuc vector by ligation (Ligate-IT, Affymetrix). Immediately following ligation, DNA was transformed and propagated in NEB stable competent cells cultured at 30°C (New England BioLabs). The resulting DNA plasmids encoding each ERS tail fused to *Gaussia* luciferase with a MANF signal peptide were sequence verified for proper ligation of the tail.

### Cell culture and transfections

Human SH-SY5Y neuroblastoma cells were maintained in growth media composed of Dulbecco’s Modified Eagle Medium (DMEM 1X + GlutaMAX, 4.5 g/L D-glucose, 110 mg/L sodium pyruvate; Invitrogen) containing 10% bovine growth serum (BGS; GE Life Sciences), 10 units/mL penicillin, and 10 µg/mL streptomycin. Cells were grown at 37°C with 5.5% CO_2_ in a humidified incubator. For reverse transfections, 5 × 10^4^ SH-SY5Y cells in 90 µL of growth media containing no antibiotics were plated in opaque TC-treated 96-well plates containing Xfect (Clontech)/DNA transfection complexes. For each well, the transfection complexes contained 200 ng plasmid DNA of unique GLuc-ERS tail and 0.06 µL of Xfect in a final volume of 10 µL. After 28-29h, a full media exchange into growth media containing 1.5% BGS was performed. Thapsigargin (Tg; Sigma) treatments began 16-17 h after the media exchange and lasted for 8 h. At the end of each experiment, media was removed from the wells and cells were lysed directly in the plate and used for luciferase assay. Cells were rinsed with 1x PBS and lysed with 75 µL of lysis buffer (50 mM Tris (pH 7.5), 150 mM NaCl, 1% NP40, 1x protease inhibitors). For all experiments, vehicle controls were used at a concentration equivalent to the Tg treatment.

### Gaussia luciferase secretion assay

Assays for *Gaussia* luciferase activity (GLuc) were performed as previously described (Henderson et al., 2014, Henderson et al., 2015). The GLuc substrate was PBS containing 10 µM coelenterazine (Regis Technologies). Coelenterazine stock solutions were prepared at 20 mM in acidified methanol (10 µL of 10 N HCl per 1 mL of methanol) and stored at -80°C as single use aliquots. For luciferase secretion assays, 5 μL of cell culture medium from each well was transferred to a new opaque walled plate. Luciferase levels were determined using a plate reader with an injector setup (BioTek Synergy II) that allowed samples to be read directly after substrate injection. 100 µL of substrate was injected to the well containing cell culture medium or lysates. For lysate readings, 100 µL of GLuc substrate was injected directly into the tissue culture plate already containing 75 µL lysis buffer. Samples were read at 25°C with a sensitivity of 100 and a 0.5 second integration time with a 5 second delay following coelenterazine substrate injection.

### KDELR2 overexpression

Lentivirus encoding Myc-FLAG tagged KDELR2 was previously described (Henderson et al., 2013). Lentiviral vectors expressing KDELR2 or a control empty vector were titered using the Lenti-X p24 rapid titer kit (Takara). 8 × 10^5^ SH-SY5Y in 2 mL growth media were plated in a 6-well plate. 24 h after plating cells were transduced with LV-KDELR2 or LV-Empty (multiplicity of infection=2). After a 48 h incubation, cells were re-plated for reverse transfections as described above.

### Machine learning based prediction model

We built recurrent neural network (RNN) models to predict Tg-induced stress response from the amino-acid sequences of last seven residues of proteins (C-terminus tails), using the TensorFlow framework (version 1.7). Each seven-residue sequence was encoded by a 7 by 20 matrix using the one-hot method, i.e., for each residue, only one element corresponding to the residue type was set to be 1 and the rest were 0 with the entire feature length of 20 representing the possible natural amino acids.

The training data was obtained from previously published results (Trychta et al., 2018a), which include 95 seven-residue tail sequences (the first batch). The cost function in the modeling process was the mean-squared error between the predicted and the true values. Based on the optimization of this function using the training data set, we chose the following setup for our models. Long Short-Term Memory network (LSTM) was chosen for our task, so that the long-term dependencies in the sequences can be captured. Only one layer of LSTM was used and the size of the hidden state of an LSTM unit was set to be 64. The LSTM layer was connected to a fully connected layer, followed by one single rectified linear unit (RELU) node to produce the output. We used the Adam algorithm to minimize the cost function with the learning rate of 0.001. The hyperparameters here were decided by 10-fold cross validation. To avoid overfitting, the network was trained with a drop-out rate 0.2, i.e. output from the 20% of the randomly chosen neurons was ignored, with 3000 iterations and during the prediction all the parameters were used.

To eliminate the stochastic element of the RNN prediction, we built multiple models to evaluate how many models were needed for the estimations to converge. Thus, for each of 64 independently built RNN models, we first calculated the Pearson correlation coefficient between the prediction and the experimental data (Figure S1A). Then for each number of models, we carried out random selection of the models from this pool (one hundred times bootstrapping) and calculated the average and the standard deviation of the Pearson R. Our results show that the slope of the standard error curve is < 0.0003 when using more than 15 models, which we define as convergence (Figure S1B). Therefore, we chose to use the average of the predictions from 32 models in our study.

### Assembling a validation tail batch from artificial sequences and human proteome

We obtained the high-frequency residue types for each position of the first batch 95 tail sequences and generated all permutations. A total of 463,736 sequences were generated and then sorted by their predicted Tg-induced stress response using our RNN models.

The entire human proteome (GRCh38.p12) consisting of 113,620 sequences was downloaded from NCBI (http://www.ncbi.nlm.nih.gov). We updated this set against the daily updating RefSeq (until 02/28/2019). For each entry in the resulting dataset, the C-terminal seven residues were extracted, and a final set of filtering was performed by removing sequences containing unnatural residues. A final set of 112,477 human protein C-terminus tails were retained, which included 31,872 unique C-terminus tails. The Tg-induced stress response prediction was then carried out for each tail using our RNN models.

We then heuristically selected 104 C-terminal seven-residue sequences, based on the following considerations: the predicted strength of responses, the novelty of the corresponding human proteins compared to the first batch, and the synthetic feasibility. The resulting batch (second batch) includes 48 artificial sequences and 56 human sequences with 65 sequences predicted to have strong responses and 39 to have weak responses.

### WebLogo and similarity scores

The sequence logo shown in Figure 2 was created using WebLogo3.6.0 (http://weblogo.threeplusone.com) to graphically represent the probability of an amino acid residue type being present at each position by the height of each symbol. The similarity scores were calculated by an in-house script using the Point Accepted Mutation (PAM)250 matrix. For each pair of aligned residues, the substitution scores were looked up from the matrix; then the similarity score for each pair of compared peptides were obtained by adding the substitution scores of -1 to -7 positions together.

### Molecular docking

The selected tail sequences were first drawn with Chemdraw (version 19.1), and then constructed with the Ligprep module of the Schrodinger (version 2019-4) suit using the OPLS3e forcefield (Roos et al., 2019). The “AEKDEL” of “TAEKDEL” was obtained from the KDELR2 crystal structure (PDB ID 6I6H), and we added “T” to the N-terminus of the peptide. The original KDELR2 structure (PDB ID 6I6H) was first processed by the protein preparation wizard module of the Schrodinger suit, and was then minimized with the OPLS3e forcefield.

We performed docking for the selected peptides with the extended sampling of the induced fit protocol (Sherman et al., 2006) implemented in Schrodinger Suite. The docking box center was determined by the center of mass of the peptide AEKDEL bound in the KDELR2 structure, and the box size was set to be 25 Å. By refining the residues within 5.0 Å of the ligand poses, the peptides were docked into the prepared KDELR2 model with the last residue “L” restrained at the original position found in the crystal structure.

### Molecular dynamic simulations

With the protein prepared as above, we also kept the crystal water inside the protein and protonated HIS 12 to HIP. The POPC lipid and water was added through Desmond setup. We kept the system charge as neutral by adding ions and the salt concentration of the system was 150 mM. The prepared system contained around 77,000 atoms, which includes ∼14,000 water molecules and 221 lipid molecules. For the peptides LIGSLEL and CIHSPDL, we chose the best docking pose as the starting position of the MD simulations. For TAEKDEL, we adopted the pose found in the crystal structure as the starting position. The MD simulations were carried out with Desmond Molecular Dynamics System (version 6.1; D.E. Shaw Research, New York, NY) using the OPLS3 force field and SPC water model. The system was first relaxed with Brownie Dynamic NVT condition with all solute heavy atoms restrained for 100 ps, and then the systems were equilibrated with NPT for 30 ns. After the equilibration, we collected two 1200 ns MD trajectories for each receptor-peptide complex system. The backbone of the transmembrane helices was restrained throughout the simulations.

## Results

### Single amino acid residue changes affect ERS function

In order to examine how changes in C-terminal ERS affect protein secretion in response to ER calcium depletion, we used a similar approach to Raykhel et al. and created a set of GLuc constructs that were identical except for their seven-residue ERS (Raykhel et al., 2007, Alanen et al., 2011). Using thapsigargin (Tg), a selective inhibitor of the sarcoplasmic endoplasmic reticulum ATPase (SERCA), we depleted ER calcium and showed that C-terminal sequences beyond the canonical KDEL motif conferred secretion in response to ER calcium depletion (Thastrup et al., 1990, Trychta et al., 2018a). To explore which permutations within the canonical KDEL motif affect the ER exodosis phenotype, we used TAEKDEL, the ERS of endoplasmin/Grp94 and BiP/Grp78, as a template to carry out an alanine scanning mutagenesis study. Our results revealed that an alanine substitution in the -1 or -2 position drastically reduced Tg-induced secretion when compared to TAEKDEL, substitutions in the -5 and -7 positions also reduced the secretion, compared to that of TAEKDEL (Figure 1). Alanen et al. found that the residues upstream of the C-terminal KDEL were important for determining ER localization of proteins and we show that the -5 and -7 positions play a role in determining whether a protein is secreted in response to ER calcium depletion (Alanen et al., 2011). These data support that ERS may play a role in regulating both protein retention (ER localization) and protein secretion. We show that changing a single residue within an ERS has the potential to change ER resident protein secretion caused by ER calcium depletion. The discovery that the amino acid composition of an ERS can affect its ability to undergo ER exodosis has ramifications for understanding how ER proteostasis changes in times of cellular stress.

**Figure 1.**
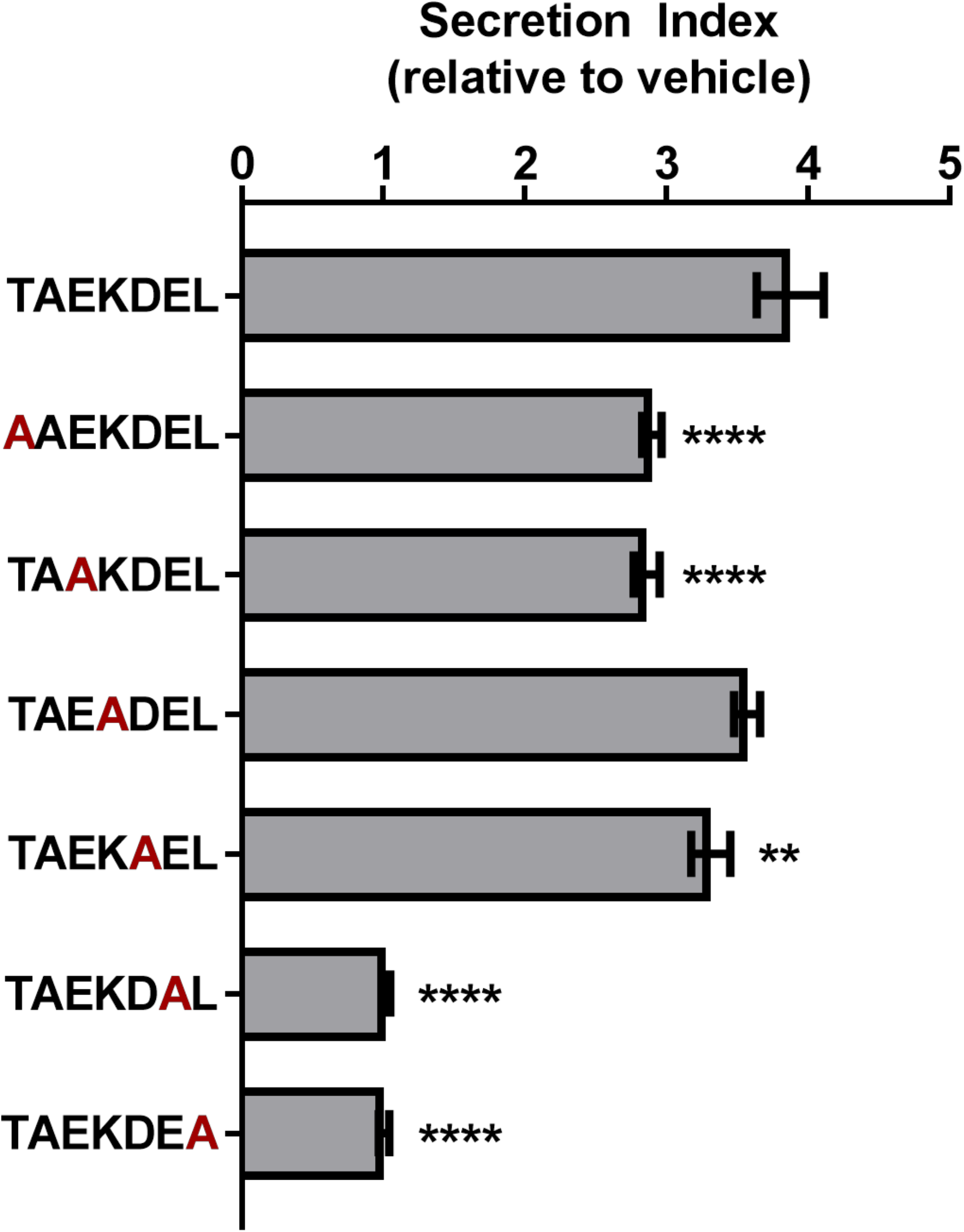
Alanine scanning mutagenesis of GLuc-TAEKDEL. Fold change in secretion of GLuc fusion proteins with TAEKDEL-like tails from transiently transfected SH-SY5Y treated with 200nM Tg for 8 h (mean ± SEM, n=36 wells, one-way ANOVA with Dunnett’s multiple comparisons, ****p<0.0001, **p<0.01).

**Figure 2.**
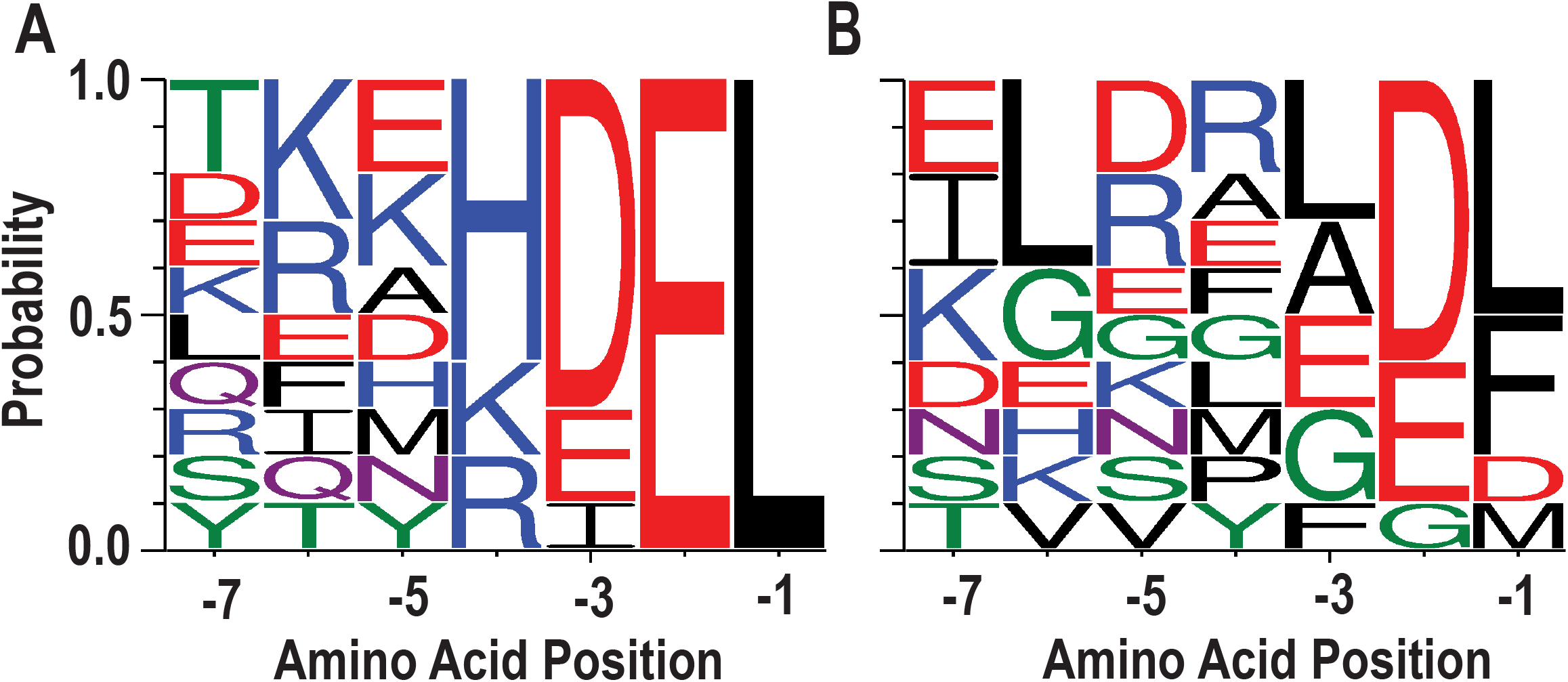
Sequences divergent from KDEL respond to ER calcium depletion. (**A**) WebLogo depicting ten peptide sequences with highest Tg response. **(B)** WebLogo depicting ten peptide sequences with lowest Tg response. For WebLogo, we used the parameters of data type “-D” as fasta, sequence type “-A” as protein, color scheme as “chemistry”, unit as “probability”.

### Non-canonical ERS undergo exodosis

Previously, we evaluated 95 different C-terminal tails from human proteins in a cell culture model of ER exodosis. We attached each seven-residue tail to the C-terminus of GLuc and monitored Tg-induced secretion responses, and found that “KKKKDEL” represented the most dominant residues at each position in the C-termini (Trychta et al., 2018a). However, when parsing the data to look at the tail sequences that result in the greatest Tg-induced secretion (top 10, Figure 2A), it is difficult to identify a consensus sequence motif that would predict secretion responses, i.e., the tails resulting in a strong response do not have a clearly dominant residue type (or residue set with similar physicochemical properties) at each position that would clearly differentiate them from the tails resulting in low responses (bottom 10, Figure 2B). For example, the negative charged residue Glu appears in -2 and -3 positions in both Figure 2A and 2B. In addition, while Leu is the only residue type appearing in the -1 position among the top 10 tail sequences, it is also the most probable residue in the bottom 10 tail sequences.

### A machine learning algorithm can predict putative ERS

Given that the canonical KDEL primary sequence motif cannot reliably predict an ERS, we used a machine learning-based approach to identify potential ERS and their connection to exodosis caused by ER calcium depletion. Using our previously published results of 7-residue tail sequences as the training data (first batch of 95) (Trychta et al., 2018a), we created recurrent neural network (RNN) models with backpropagation and long short-term memory (LSTM) algorithm (Hochreiter and Schmidhuber, 1997). The models were then used to predict 7-residue tail sequences for their responsiveness to Tg treatment (i.e. ER calcium depletion). Based on the predictions by our models, we selected 104 sequences (second batch) for experimental validation. The selected sequences included both those from human proteins and artificial ones that do not exist in the human proteome. In addition to selecting sequences predicted to have strong Tg responses, we also included some sequences predicted to have weak Tg responses to validate our models. The selected ERS were appended to the C-terminus of GLuc and used for validation in our experimental model of ER calcium depletion.

Comparing the predicted Tg responses (referred to as RNN scores) and the experimentally determined Tg responses, moderate positive correlations were found based on either the response values or their relative rankings, with correlation coefficients of 0.70 and 0.69, respectively (Figure 3A-B). Interestingly, a few artificial sequences that were predicted to have high Tg-responses were indeed found to experimentally exhibit higher responses than any of the human sequences tested in the second batch (Figure 3B). In addition, our findings further demonstrate that the primary KDEL sequence does not include adequate information to predict ER protein secretion. For example, in both the first and second batches of data, the positively charged Arg and Lys appear to be dominant but the oppositely charged Asp and Glu can be tolerated. This can be observed for KHLEDEL which had an RNN score of 4.1 ± 0.1 and an experimentally determined Tg response of 5.2.

**Figure 3.**
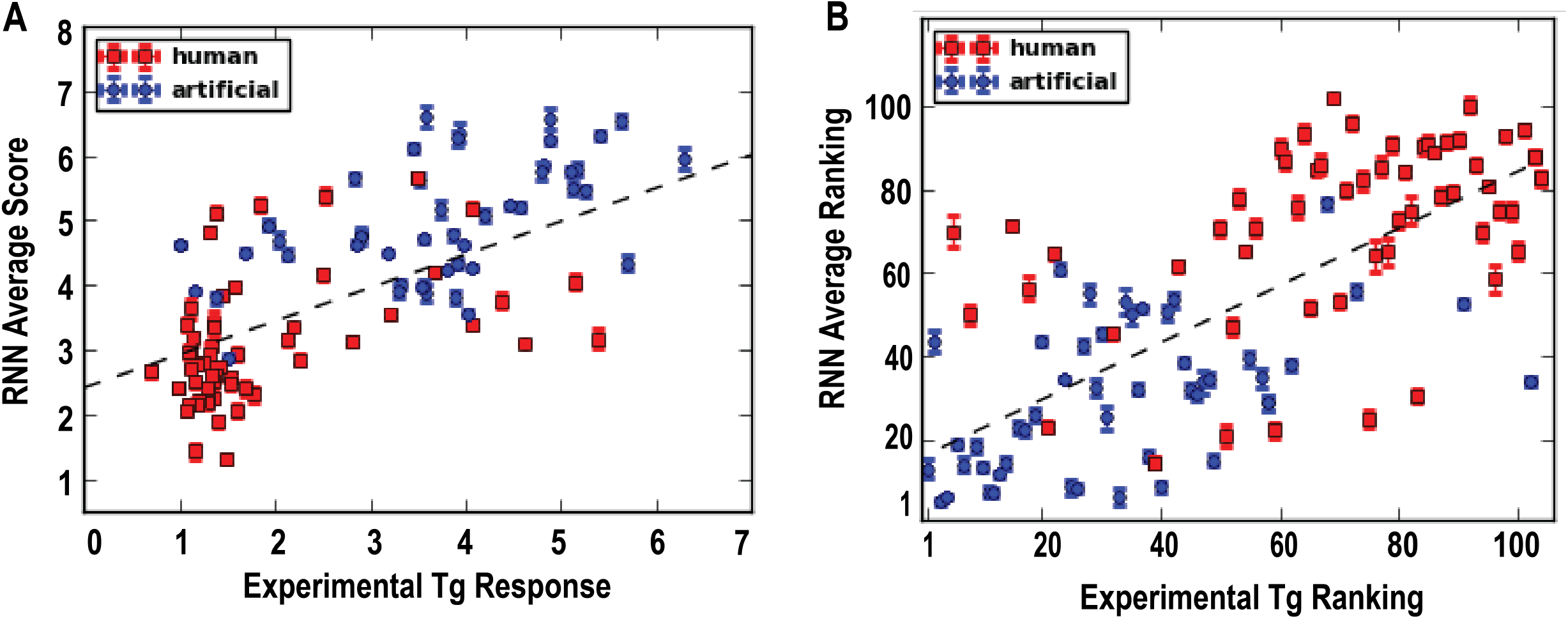
Machine learning predicts ERS response to ER calcium depletion. Recurrent neural network (RNN) scores were obtained by feeding the second batch data to the RNN models. The prediction ranking was based on the sorting of the RNN scores by the descending order. **(A)** Correlation between predicted RNN average scores and experimentally determined Tg responses (correlation coefficient: 0.70). **(B)** Correlation between the ranking of the RNN scores and the ranking of experimentally determined Tg response (correlation coefficient: 0.69).

### Non-KDEL ERS form alternative interactions with KDEL receptor

To understand the molecular recognition between the C-terminal ERS and KDEL receptors, we performed molecular modeling studies of selected ERS. A recent crystal structure of chicken KDEL receptor 2 (KDELR2) demonstrated how the canonical ERS TAEKDEL binds to KDELR2 (Bräuer et al., 2019). In the crystal structure, the deeply buried -1Leu and -2Glu of the bound peptide make extensive interactions with the receptor, and directly contact 10 and 6 residues respectively; in comparison, -3Asp and -4Lys interact with 3 and 4 residues, respectively (Figure 4A, Supplementary Figure S3A). To investigate these varied extents of interactions in the recognition of the ERS by KDEL receptors, we first built a simulation system of KDELR2 bound with TAEKDEL and embedded in explicit water-lipid bilayer environment, and carried out molecular dynamics (MD) simulations (see Methods). In our simulations, we observed that the TAEKDEL peptide largely retained the pose found in the crystal structure (Figure 4B, Supplementary Figure S3B). However, the position of the -3Asp sidechain in the crystal structure appears to be in a high-energy state, which is stabilized only by its interaction with Arg169. In our simulations, while the χ1 rotamer of -3Asp can be in the gauche+ (∼+60°) position for much time as in the structure, it frequently visited the gauche-(∼-60°) position (Supplementary Figure S2A), a more favored position for Asp (Lovell et al., 2000). In comparison, the χ1 of -2Glu residue was persistently in its favored gauche-rotamer throughout our simulations (Supplementary Figure S2A). When χ1 of -3Asp is in gauche+, it forms an ionic interaction with Arg169; when it is in gauche-, it loses this interaction and becomes dynamic. In comparison, -2Glu persistently forms a salt bridge with Arg169 (Supplementary Figure S2).

**Figure 4.**
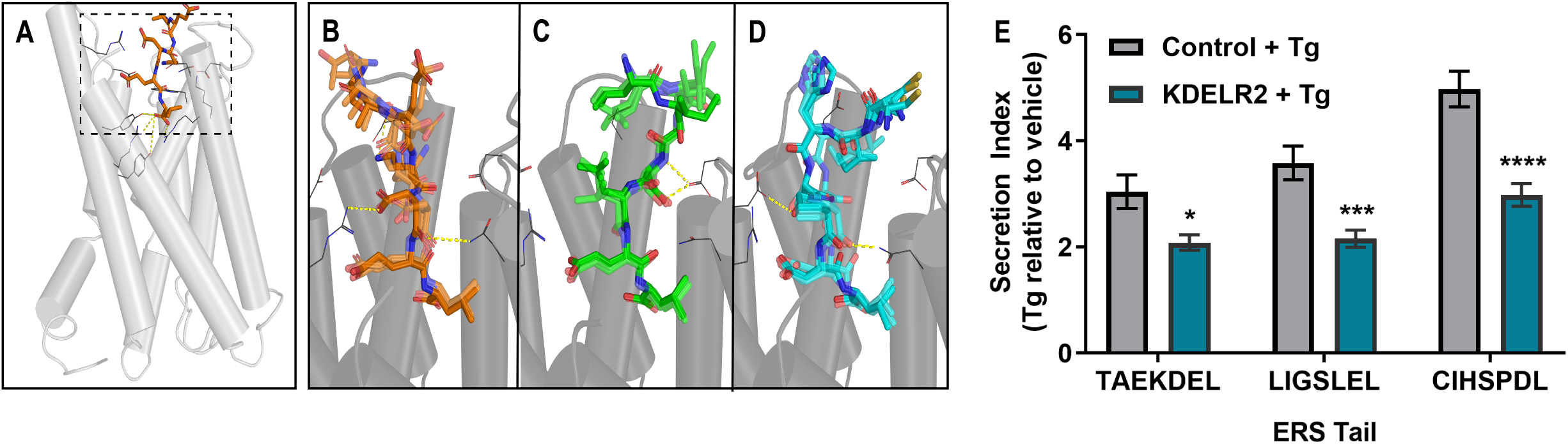
Sequences interact with the KDEL receptor. (A) Overview of KDEL receptor 2 crystallized structure (PDB ID: 6I6H). The dashed box highlights the binding site. The crystal peptide AEKDEL is depicted in orange. **(B)** Representative frames selected from the TAEKDEL MD simulation. The peptide TAEKDEL is shown in orange, and the receptor is shown in gray. Yellow dashed lines indicate where interactions between the peptide and the KDEL receptor occur. **(C)** Representative frames selected from the LIGSLEL MD simulation. Peptide LIGSLEL was shown in green. **(D)** Representative frames selected from the CIHSPDL MD simulation. Peptide CIHSPDL is shown in blue. **(E)** Fold change in secretion of GLuc-ERS proteins from transiently transfected SH-SY5Y cells overexpressing KDELR2 or control following treatment with 200 nM Tg for 8 h (n=14, two-way ANOVA with Sidak’s multiple comparisons, **p<0.01, ****p<0.0001).

This finding of the relatively weak interaction between -3Asp and KDELR2 suggests that replacement of Asp at this position can be tolerated for other ERS. We found that alanine substitution at -3Asp had only a modest effect on basal secretion (from data in Figure 1). When we calculated the similarity scores of our experimentally verified ERS against KKKKDEL, the most probable amino acid motif derived from our previously identified WebLogo sequence (see also Methods), we found two ERS that are most divergent from KKKKDEL, LIGSLEL and CIHSPDL, do not have Asp at the -3 position. Compared to the RNN score of TAEKDEL, 4.99, those for LIGSLEL are CIHSPDL are 3.26 and 4.56, respectively. To characterize how these two peptides interact with KDELR2, we docked them in KDELR2, similarly built the simulation systems as with KDELR2/TAEKDEL, and carried out MD simulations (see Methods). In our simulations, the -3Leu of LIGSLEL does not make any interaction with the receptor, while -3Pro of CIHSPDL can stably interact with Arg5, Asn60, and Arg169. Even though neither peptide has the -4Lys seen in KKKKDEL, for LIGSLEL, both the sidechain and backbone of -4Ser persistently interact with Asp50 (Figure 4C, Supplementary Figure S3C), while the -4Ser of CIHSPDL interacts with Asp177 keeping the peptide in a stable conformation (Figure 4D, Supplementary Figure S3D).

The -5, -6, and -7 position residues of the peptides have fewer interactions with the receptor compared to the -1 to -4 residues because they are located at the opening of the binding pocket. However, interestingly for different peptides, residues in the -5, -6, or -7 positions interacted with the different residues of the receptor. Specifically, -7Thr of TAEKDEL interacts with Asp112 of the receptor, while the -6Ile of LIGSLEL and ILE56 of the receptor forms a hydrophobic interaction. For CIHSPDL, -6ILE interacts with four residues, in particular, the hydrophobic interaction with Leu116 of the receptor appears to be a stabilizing force by interacting with -6L (Supplementary Figure S3).

Despite their divergence from the canonical KDEL sequence, both peptides can be reasonably adopted in the KDEL receptor binding site, suggesting that KDEL receptors can interact with a variety of ERS tails and that Tg-induced ERS protein secretion is not due to a lack of interaction with KDEL receptors. KDEL receptors normally function to return escaped ER resident proteins from the Golgi to the ER lumen, but following ER calcium depletion KDEL receptors are overwhelmed by the mass efflux of proteins from the ER resulting in increased ERS protein secretion (Trychta et al., 2018a). Overexpression of KDELR2 can attenuate the Tg-induced secretion of GLuc-TAEKDEL, as well as that of GLuc-LIGSLEL and GLuc-CIHSPDL (Figure 4E). The increased secretion of LIGSLEL and CIHSPDL constructs following Tg treatment and their predicted interaction with KDELR2 indicate they act in a manner consistent with an ERS secretion phenotype. These data suggest that proteins SFTPB and CNTN2, which contain the predicted ERS LIGSLEL and CIHSPDL, respectively, would be secreted in conditions associated with ER calcium depletion and, in fact, increased secretion of these proteins has been associated with disease (Supplementary Table S2).

## Discussion

Here we investigate C-terminal ERS sequences and assess how their amino acid sequence can predict protein secretion following ER calcium depletion. Our recent work suggests that ER calcium depletion causes ERS-containing proteins to be secreted from the cell (Trychta et al., 2018a). The ability to predict whether a protein would be secreted in response to reductions of ER luminal calcium could assist in our understanding of the role of ER luminal proteins in pathological conditions. A loss of essential luminal proteins, i.e., exodosis, could negatively impact ER functioning, while the presence of these normally ER luminal proteins outside of the cell could also change the extracellular environment. A gain of function outside of the cell could be detrimental if the secreted proteins contribute to inflammation or lead to aberrant activation of signaling pathways, but the extracellular localization of some ERS-containing proteins could have beneficial effects. For example, in rheumatoid arthritis BiP at the cell surface can stimulate tumor necrosis factor (TNF) production to promote inflammation, but BiP can also act as an antigen to stimulate CD8-positive T cells leading to increased production of the anti-inflammatory cytokine IL-10 (Bodman-Smith et al., 2003, Lu et al., 2010). Evaluating whether a protein has a putative ERS may provide mechanistic and etiological understanding of related diseases.

Published data also supports that ERS-containing proteins can saturate KDEL receptors when they leave the ER en masse (Trychta et al., 2018a). In the current molecular modeling study we show that ERS divergent from the canonical KDEL motif can be reasonably accommodated in the KDEL receptor binding site, which is consistent with the observation that overexpression of KDELR2 can attenuate the secretion of proteins with these divergent ERS (Fig. 4). This evidence of an interaction with KDEL receptors warrants further study to gain a complete picture of the KDEL receptor retrieval pathway and how ERS differences or KDEL receptor isoforms may influence protein trafficking. Furthermore, the understanding of the presence and action of KDEL receptors on the plasma membrane may provide insight into the mechanism of action of ERS containing proteins that are secreted from the cell following ER calcium depletion (Jia et al., 2020).

The implications of our machine learning algorithm go beyond predicting whether a protein is secreted following ER calcium depletion based on its C-terminal ERS or whether an ERS can dock to KDEL receptors. In this study, we used a Tg-based model of ER calcium depletion, but there are many disease states associated with ER calcium depletion (Mekahli et al., 2011). The ability to predict which proteins may be secreted from the cell in response to calcium depletion can help guide studies of the complex relationship between disruptions in ER homeostasis and diverse pathologies. Our data support that artificial sequences not present in the human proteome could be added to reporters or therapeutic proteins to achieve different degrees of protein secretion in response to calcium depletion. Given our model also predicts proteins that will have an increased extracellular abundance in diseases associated with ER calcium depletion, the development of new biomarkers is feasible. Additionally, for diseases in which ERS proteins have previously been found in increased amounts in extracellular fluids, the role of ER calcium depletion and exodosis in the pathology of the disease may warrant further study. For example, the increased secretion of luciferase reporters with LIGSLEL and CIHSPDL tails suggests their corresponding proteins, SFTPB and CNTN2, respectively, would be elevated in response to ER calcium depletion. Interestingly, the mouse SFTPB, which has the tail CFQTPHL, was also shown to be increased in plasma of a cancer model (Taguchi et al., 2011). Our model predicted CFQTPHL to have an RNN score of 4.36, which is comparable to that of CIHSPDL (4.56). Even though the similarity of CFQTPHL and CIHSPDL is not high, they both have -3Pro, which was found to make extensive interactions with KDELR2 (Figure 4), while at the -4 position, Thr is expected to make similar interactions as Ser. In fact, both proteins have been shown to be elevated extracellularly in diseases (Supplementary Table S2) and may serve as biomarkers and provide insight into disease mechanism as well as therapeutic approaches to stabilizing ER calcium and preventing exodosis.

We show that ERS divergent from the prototypical sequence “KDEL” are capable of binding to the KDELR and with different interacting residues on the KDELR. The term “KDEL receptor” does not fully reflect the breadth of ligands to which it binds. Our paper helps to further illuminate the diverse ligands that can bind to the KDEL receptors and be retained within the cell but secreted upon decreases in ER calcium. Future studies examining the structural differences among the KDELR isoforms and their preferences for different ERS ligands will be necessary to better understand their role in this complex protein retrieval pathway.

Overall, we present a systematic examination of the connection between C-terminal ERS, KDEL receptors, and protein secretion and highlight the combined machine-learning and molecular modeling approaches that can predict and characterize ER resident protein secretion mediated by ER calcium depletion and recognition by the KDEL receptor.

## Acknowledgements

This work was supported by the National Institute on Drug Abuse, Intramural Research Program (Z1A DA000606 to L.S. and Z1A DA000618 to B.K.H.).

## Figure Legends

**Supplementary Table S1: ERS proteins are increased in the extracellular fluid in pathological conditions**.

**Supplementary Table S2: SFTPB and CNTN2 are increased extracellularly in disease states**.

**Supplementary Figure S1:**
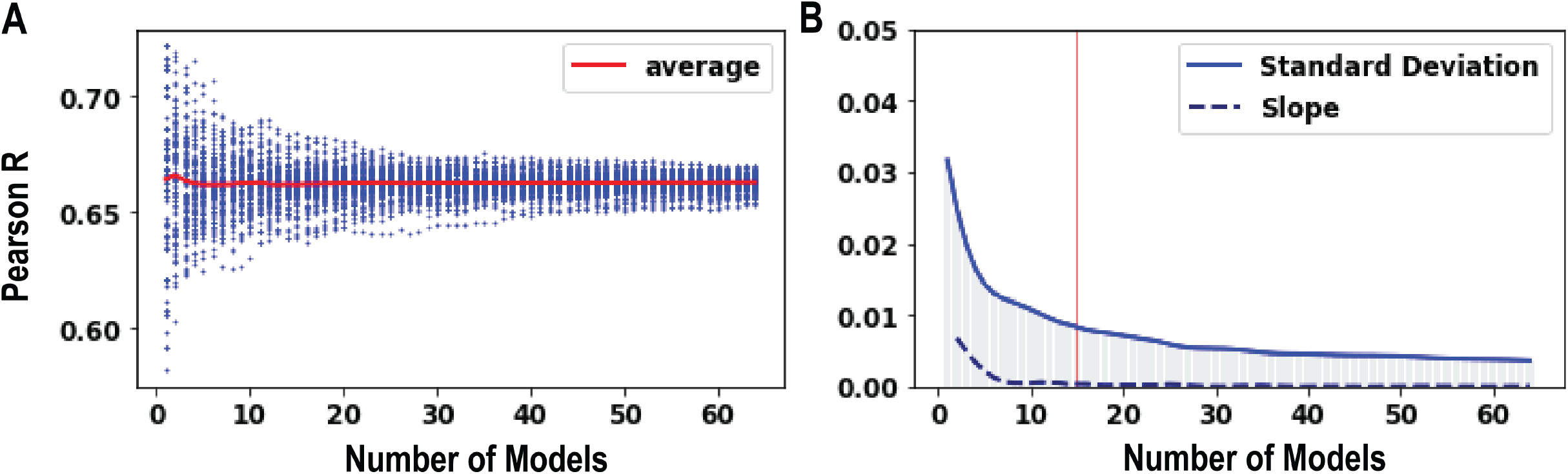
Determining the number of models needed to predict results. **(A)** With 100 times bootstrapping, more models made the correlation estimation between predicted RNN scores and experimental Tg response converge. Each single blue line represents one bootstrapping result. **(B)** The standard error of correlation decreased with an increasing number of models. After the slope reached 0.0003, the calculation was converged with the red line indicating that this convergence occurred with 15 models.

**Supplementary Figure S2:**
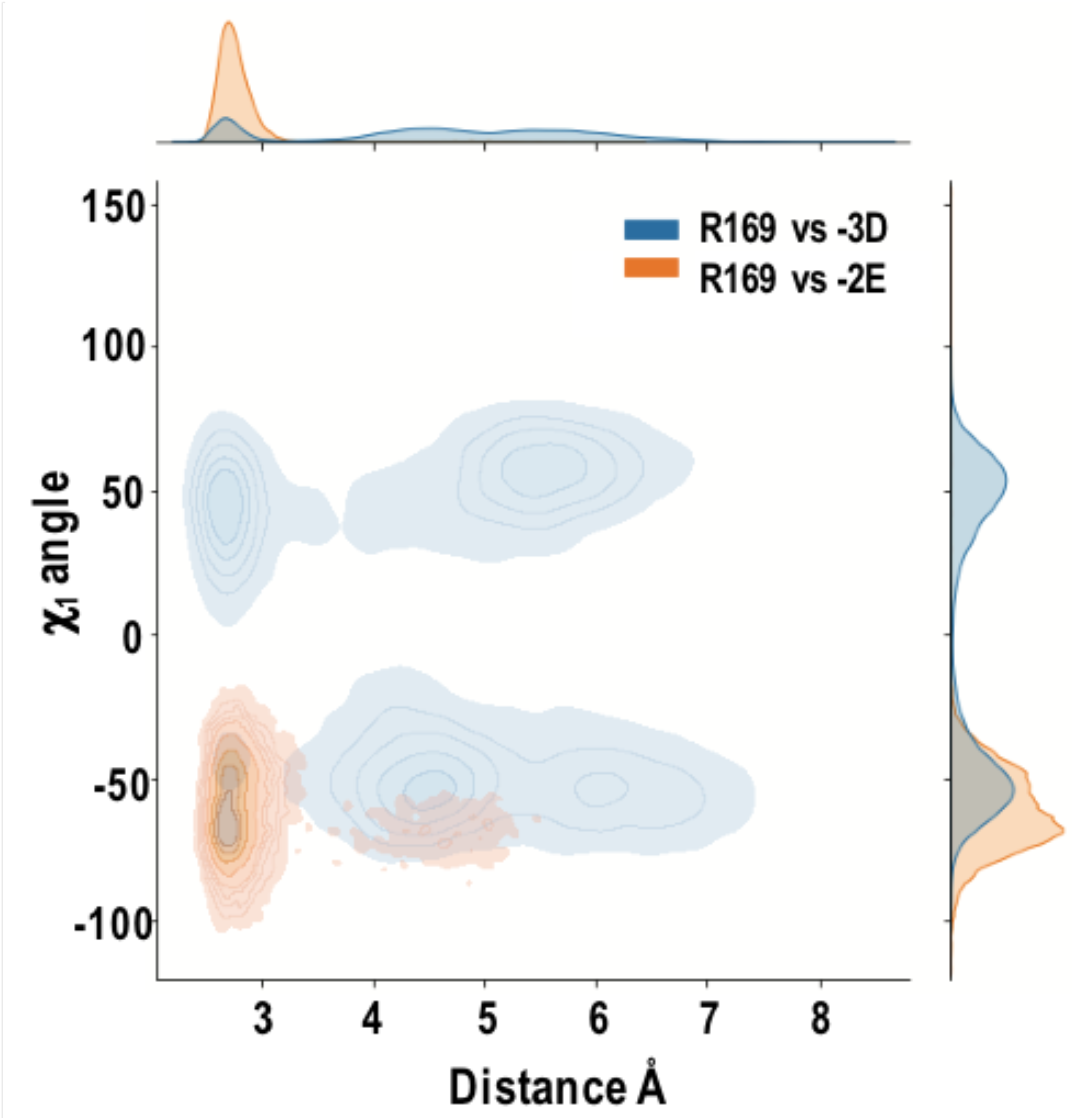
The -2 and -3 position of TAEKDEL exhibited different stability during simulated KDEL receptor docking. The x-axis indicated the distance between the peptide and the KDEL receptor, while the y-axis indicates the dihedral angles of the TAEKDEL peptide.

**Supplementary Figure S3:**
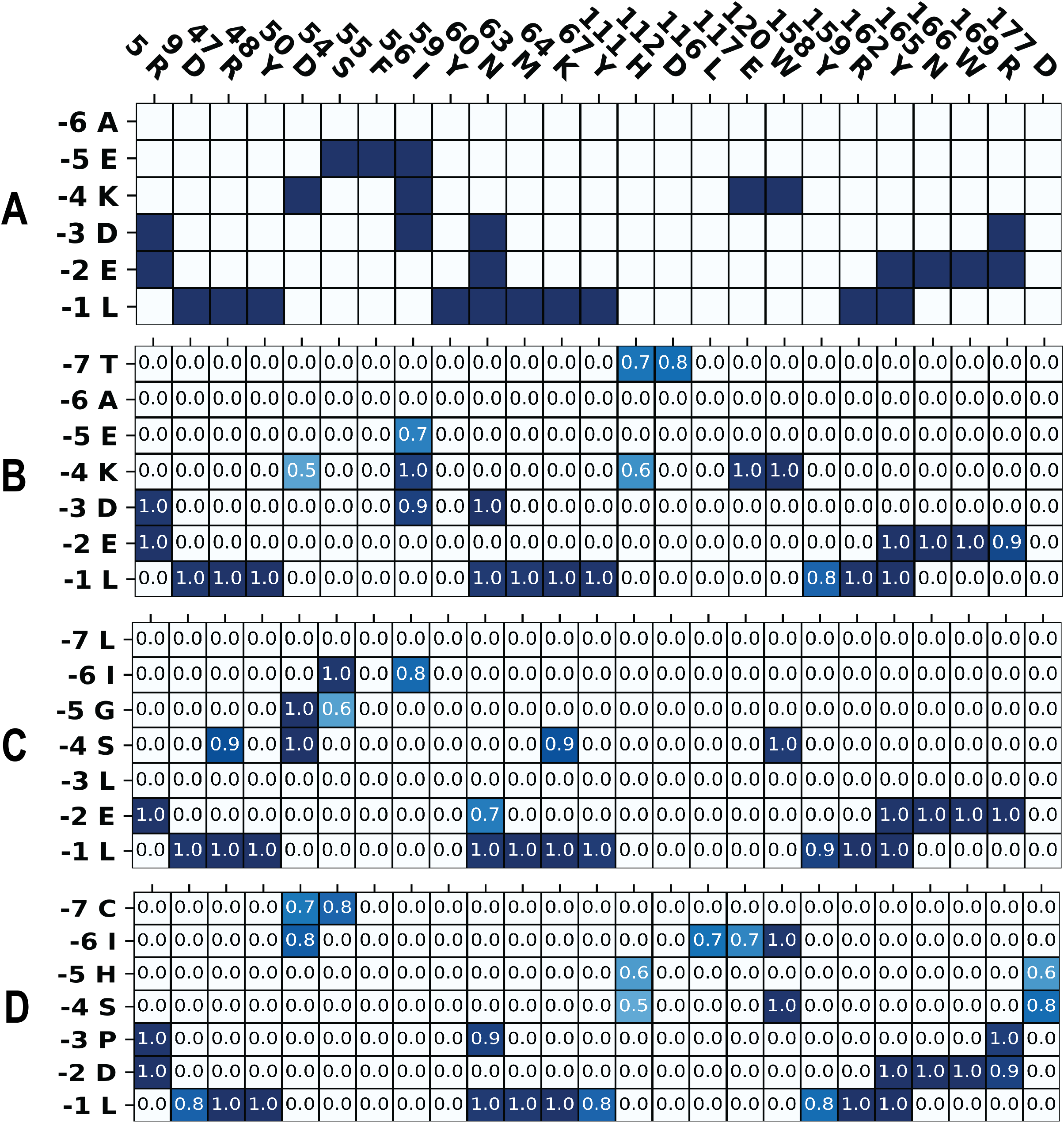
Heatmap of contact frequency between ligand and the KDEL receptor. **(A)** The interaction between crystal peptide AEKDEL and the KDEL receptor crystal structure. **(B-D)** The interactions between **(B)** TAEKDEL, **(C)** LIGSLEL, or **(D)** CIHSPDL and the KDEL receptor.

**Supplementary Table S1:**
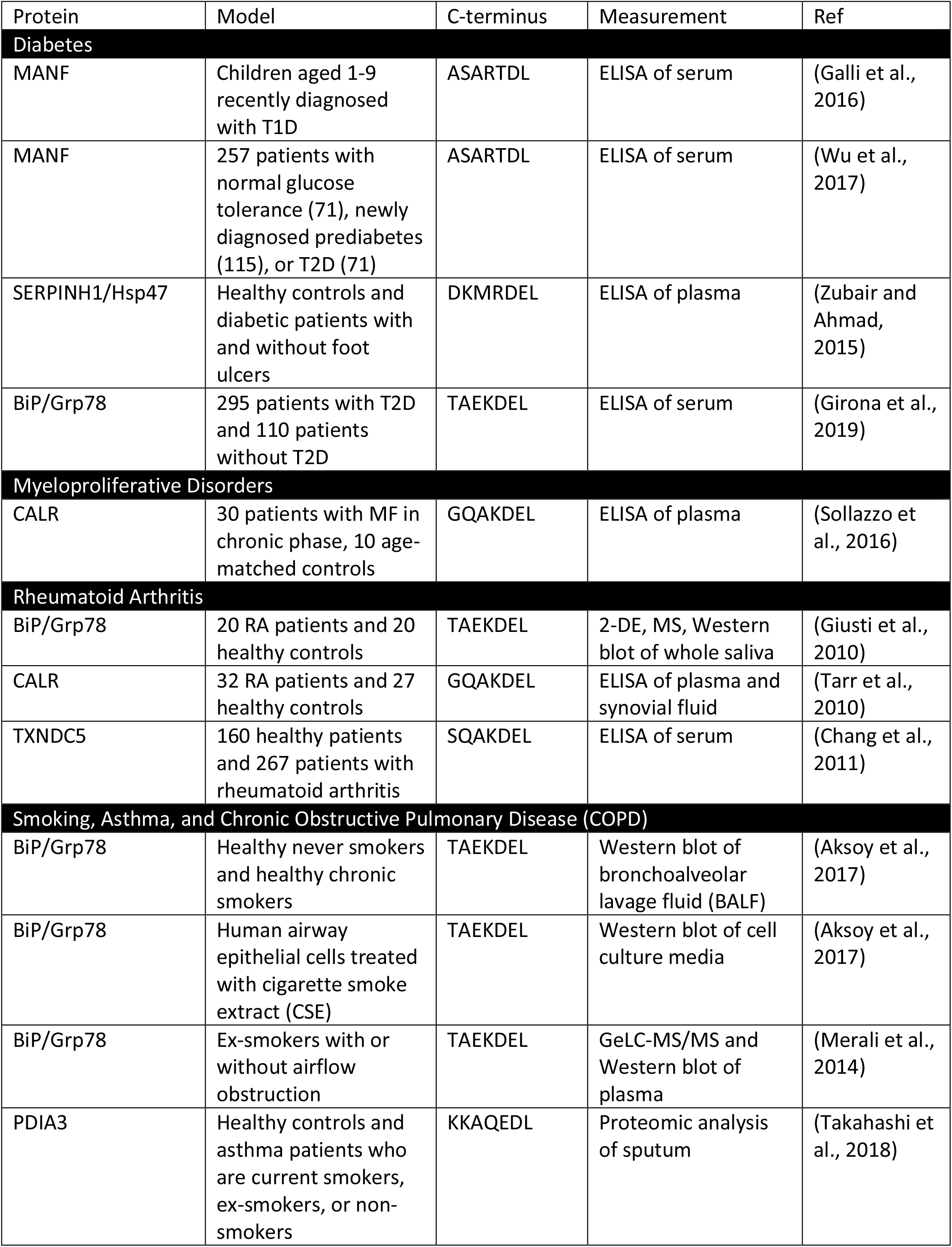
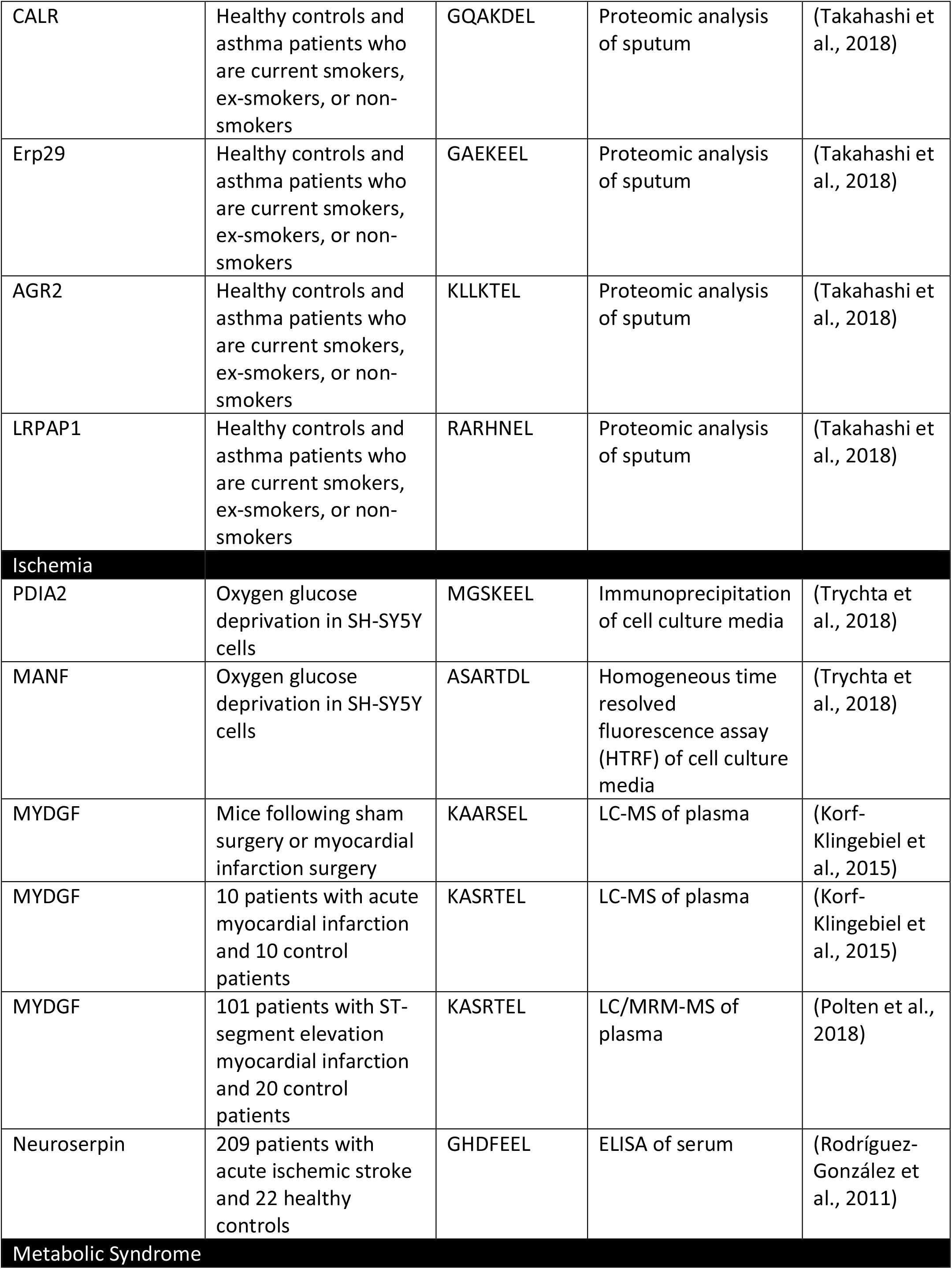
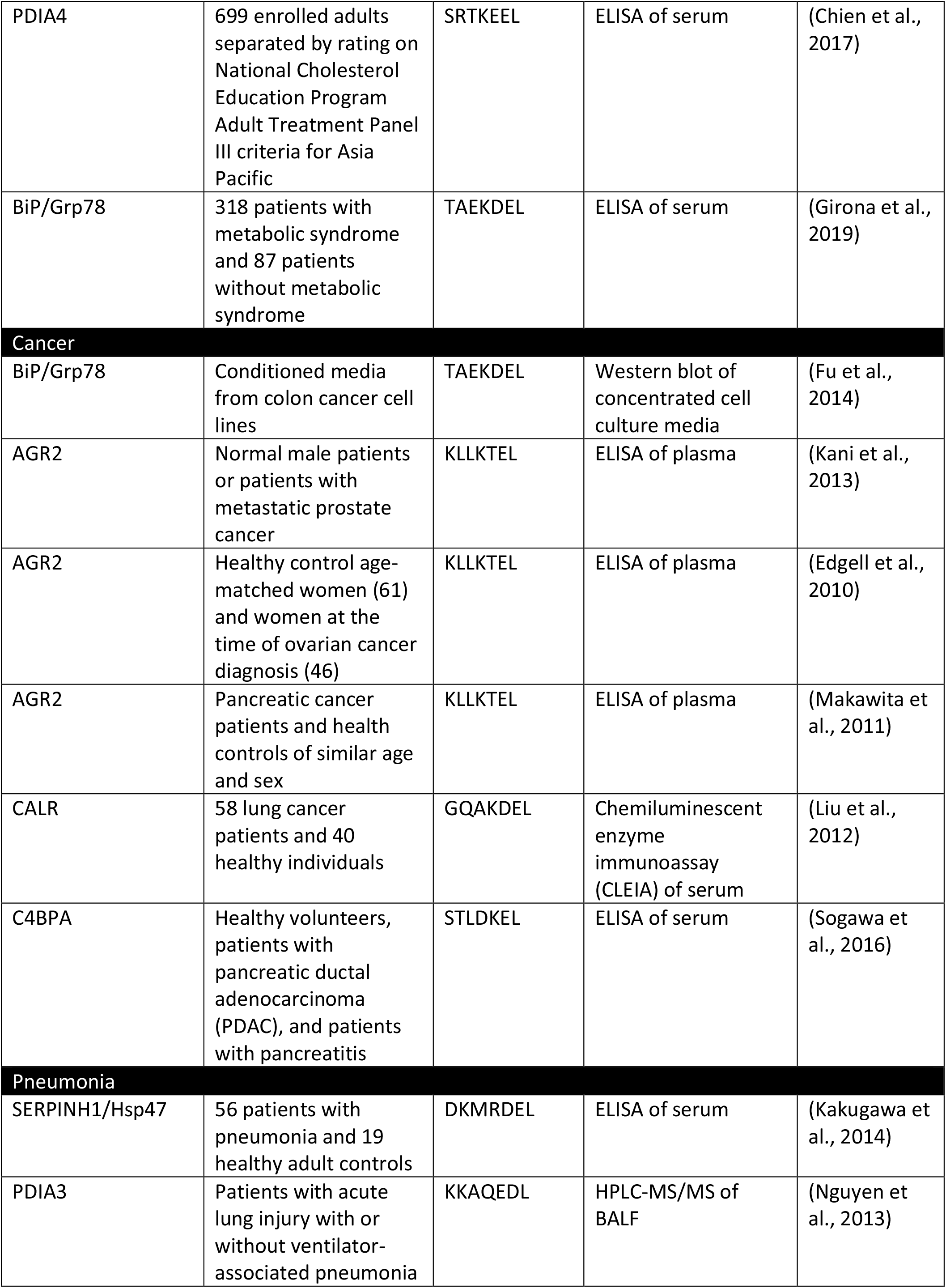
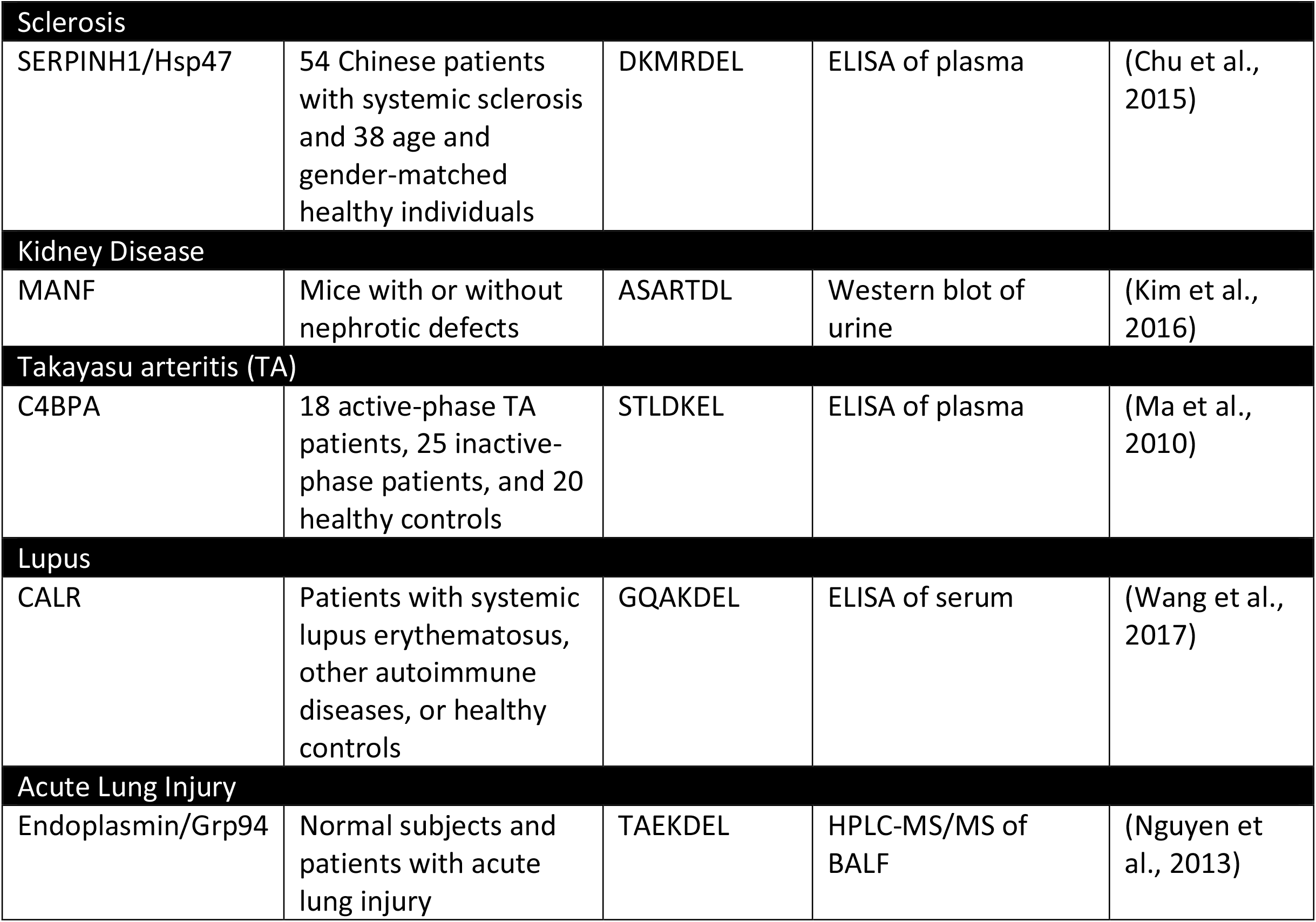
ERS proteins are increased in the extracellular fluid in disease states.

**Supplementary Table S2:** SFTPB and CNTN2 are increased extracellularly in disease states.

| Protein | Model | C-terminus | Measurement | Ref |
| --- | --- | --- | --- | --- |
| **Cancer** |  |  |  |  |
| SFTPb | Tumor bearing mice from lung adenocarcinoma models and controls | CFQTPHL | MS of plasma | (Taguchi et al., 2011) |
| SFTPb | Subjects with diagnosed non-small cell lung carcinoma and controls | CIHSPDL | ELISA of plasma | (Taguchi et al., 2011) |
| **Lung Disorders** |  |  |  |  |
| SFTPb | Patients diagnosed with acute respiratory distress syndrome or controls | CIHSPDL | ELISA of plasma | (Doyle et al., 1997) |
| SFTPb | Patients diagnosed with acute cardiogenic pulmonary edema or controls | CIHSPDL | ELISA of plasma | (Doyle et al., 1997) |
| SFTPb | Current or never smokers | CIHSPDL | ELISA of plasma | (Nguyen et al., 2011) |
| SFTPb | 15 infants with respiratory syncytial virus and 13 healthy controls | CIHSPDL | ELISA of plasma | (Wang et al., 1999) |
| SFTPb | Current smokers and non-smokers | CIHSPDL | ELISA of serum | (Robin et al., 2002) |
| SFTPb | People before and after attendance at a chlorinated pool with nitrogen trichloride gas exposure | CIHSPDL | ELISA of serum | (Carbonnelle et al., 2002) |
| **Ischemia** |  |  |  |  |
| SFTPb | Patients with a cerebral infarction or controls | CIHSPDL | ELISA of CSF | (Schob et al., 2013) |
| **Alzheimer's disease** |  |  |  |  |
| CNTN2 | 5 AD patients and 6 normal subjects | LIGSLEL | LC-MS/MS of CSF | (Yin et al., 2009) |
| **Heart disease** |  |  |  |  |
| SFTPb | 53 outpatients with chronic heart failure and 19 healthy controls | CIHSPDL | ELISA of plasma | (De Pasquale Carmine et al., 2004) |
| SFTPb | Patients with exercise-induced myocardial ischemia pre and post-exercise | CIHSPDL | ELISA of plasma | (De Pasquale et al., 2005) |

